# A colorimetric method for detecting virulent bacteriophage to *Vibrio cholerae* in fecal and environmental samples

**DOI:** 10.1101/2025.08.07.669158

**Authors:** Wensheng Luo, Diana Dinowitz, Andrew Camilli, Jason R Andrews, Kesia Esther da Silva, Amanda K Debes, Md A Sayeed, Eric J Nelson, David A Sack

**Affiliations:** Department of International Health, Johns Hopkins Bloomberg School of Public health, Baltimore, Maryland; Department of Molecular Biology & Microbiology, Tufts University School of Medicine, Boston, MA; Division of Infectious Diseases and Geographic Medicine, School of Medicine, Stanford University, Stanford, California; Departments of Environmental and Global Health and Pediatrics, Emerging Pathogens Institute, University of Florida

**Keywords:** cholera, vibrio, vibriophage, bacteriophage, phage, resazurin, *Vibrio cholerae*, environmental samples, wastewater, AC6169

## Abstract

**Background:** *V. cholerae* are often infected with vibriophages and these phages may be found associated with *V. cholerae* bacteria or may be detected independently from the bacteria, especially in environmental water samples. When detected, vibriophages serve as a surrogate for detection of *V. cholerae* and they also have an important role in the ecology of *V. cholerae*. Vibriophages can be detected using plaque assays or PCR, but these methods are time-consuming and require specialized laboratory resources.

**Methodology:** To address limitations of other methods, we developed a simple, rapid, and inexpensive colorimetric assay to detect vibriophage that can be scaled up to quickly and easily to screen a large number of samples. The assay uses resazurin and a bacterium, AC6169 that is susceptible to vibriophages ICP1, 2, and 3. Resazurin is a dye that turns color from blue to pink when added to a culture broth with growing bacteria. When a bacteria-free test sample, such as a Millipore filtered wastewater sample containing vibriophages, is added to a culture broth with AC6169, the bacteria will be lysed by the phages and will not grow, and the color of the broth will remain blue. However, if there are no phages in the sample, the bacteria will grow rapidly, and the culture broth will turn pink. We developed the assay using ICP1 spiked samples of environmental water, stool and frozen bile peptone and found it to be sensitive with a limit of detection of 4 to 40 plaque forming units/ml.

**Conclusion:** This colorimetric assay promises to provide a convenient method to detect vibriophages on a larger scale than was possible earlier to better understand their role as a surrogate for detecting V. cholerae and to better understand their role in the pathogenesis, ecology, and epidemiology of cholera.

**Author Summary:** We developed a simple and inexpensive colorimetric assay to detect virulent vibriophage in samples of wastewater and stool. The assay uses resazurin, a blue dye and a unique bacterium, AC6169, that is known to be susceptible to vibriophages. Bacteria-free samples (e.g. Millipore filtrates) suspected of having vibriophage are incubated for 2 or 3 hours in a broth with AC6169, following which resazurin is added. After a further 30-minute incubation, the color of the broth is observed. If the sample contains vibriophage, the broth will remain blue, but if there are no vibriophage, the broth will turn pink. The change in color from blue to pink is due to the metabolic reaction from the growing bacteria. When vibriophage are present, they lyse the bacterium AC6169, thus preventing the color change, but if vibriophage are not present, the bacteria multiply rapidly, and the metabolic reaction causes the broth to change color to pink. Because the assay procedure is inexpensive and simple, laboratories in cholera endemic areas should find it convenient for their epidemiologic and clinical surveillance activities.

## Introduction

Cholera, a bacterial diarrheal disease caused by *Vibrio cholerae*, is characterized by severe watery diarrhea that can quickly lead to severe dehydration and death(1). This disease continues to be a major public health problem and the World Health Organization considers cholera to be an “Ongoing Health Emergency(2).” Currently an estimated 2.9 million cases with 95,000 deaths occur each year(3). Patients with cholera are sometimes co-infected with virulent vibriophage (phage) which prey on the infecting *V. cholerae*. When these phage-infected bacteria are excreted in the patient’s stool, the phages are also shed. In areas with poor sanitation, they may be co-ingested by the next human host. Thus, infections with *V. cholerae* can occur either with phage or without phage, but phages cannot persist for a long time without the *V. cholerae* on which they prey(4).

Because the vibriophage exist in a close relationship with *V. cholerae*, the detection of the phage is an indicator that *V. cholerae* are present(5). Therefore, when vibriophages are detected in a stool sample of a patient suspected of having cholera, this may also confirm the diagnosis of cholera. Similarly, when vibriophage are identified in an environmental sample, such as wastewater, this provides evidence that *V. cholerae* is also being expelled into the environment even if the bacterium itself is not detected in the sample(6). This concept can be called diagnosis by phage proxy.

Most diagnostic efforts on cholera are directed toward detecting the *V. cholerae* bacterium either by rapid test, culture or by PCR. To supplement these efforts, we felt it is important to also explore methods to detect vibriophage simply and rapidly in patients’ stools and in environmental samples. Traditional methods for detecting phage have used plaque assays(7). When present, these plaques form on an agar plate inoculated with susceptible *V. cholerae* and the plaque allows one to identify a phage that had preyed on the target bacteria. These areas of lysis can then be harvested and can be cloned to purify and characterize the phage that has infected the bacterial cells. Using this method three different O1 El Tor-specific vibriophages have been isolated from patient’s stool samples in Bangladesh and have been named ICP1, ICP2 and ICP3(8). Subsequently, these phages were sequenced, and PCR methods developed to detect phage in stool samples, including dried fecal samples on filter paper (unpublished, E, Nelson). Using plaque assays, a phage closely related to ICP1 was detected in the Democratic Republic of the Congo(9).

While plaque assays and PCR can be carried out to detect vibriophage, these require time, technical laboratory skills and resources. We felt a colorimetric method using the dye resazurin might be adapted to screen fecal and environmental samples more quickly and inexpensively. Such an assay could facilitate large-scale screening efforts in both clinical and environmental settings, ultimately improving cholera surveillance and outbreak response. Similar methods have been used to detect or study phage-specific for *E. coli* and *Salmonellae*(10-13). Resazurin, a blue dye, is converted into pink, fluorescent resorufin in the presence of metabolically active cells. NADPH dehydrogenase from the multiplying bacteria is thought responsible for the conversion of resazurin to resorufin. This conversion can be detected through the visual observation of its pink color. Resazurin has also been used for determining antibiotic susceptibility of bacteria(14, 15) and fungi and for screening natural materials for their inhibitory ability against bacteria(16). We hypothesized that if vibriophages are present in a test sample consisting of bacterial-free filtered feces or environmental water, it will lyse a pan-sensitive *V. cholerae* test strain. Thus, in the presence of resazurin, the susceptible bacteria in a culture broth mixed with a test sample with phages will fail to grow and the culture broth will remain blue. However, if the test sample does not have phages, the bacteria will grow rapidly, and the broth will turn pink. Ideally, this change would be apparent to the naked eye; however, the color change could also be objectively determined by measuring the OD_600_ of the culture supernate. OD_600_ is the appropriate wavelength because this is the peak absorbance for the blue dye, compared to the pink dye with a peak of 570 nm(17). Although ICP1, 2 and 3 were identified in Bangladesh, this manuscript focuses only on ICP1 since it has been identified in Africa where there are multiple ongoing cholera outbreaks(9, 18).

## Methods

A series of experiments were carried out to explore methods appropriate for phage detection. These methods are modified from methods used to detect phage for *Salmonella* Typhi(13). The experiments used varying concentrations of ICP1 which were spiked into samples of environmental water, stool, or bile peptone broth (BP) to determine the sensitivity of detection in these samples.

### Materials and preparation of the materials

Environmental water was obtained from the northern area of the Chesapeake Bay. Bile peptone (BP) was prepared according to an established recipe of 20g Ox Bile (Millipore-Sigma #70168-100g), 5g Dextrose(Fisher Chemicals #D16-500), 8g Sodium phosphate dibasic (Santa Cruz #c-203277), 10g Peptone from gelatin (Millipore-Sigma,#107284.100), and 2g potassium dihydrogen phosphate (Fisher # P285-500) in one liter with water and autoclaved for 15 minutes at 121°C). Vials of BP were spiked with varying concentrations of phage and were then frozen (−80°C) until tested. Fresh dog stools were diluted 1:5 in PBS to simulate a diarrheal stool.

Vibriophage ICP1_2011_A^7^ and phage indicator strain *V. cholerae* strain AC6169 were used in this study. AC6169 is a genetically engineered, non-toxigenic, antibiotic-sensitive, Biosafety Level 1 (BSL-1) derivative of the O1 serogroup, El Tor biotype strain E7946. This strain was constructed using the suicide plasmid pCVD442 to mediate several allelic exchange events as described(19). First, the CTX prophage and flanking RS1 and TLC mobile elements were deleted (fusion junction sequence 5’-GTGGAGTTCTTTTTT/GCACTGAGGTATTTT-3’). Second, the K139 prophage and its *attB* site were deleted (fusion junction sequence 5’-TGGGGGTAAAACGC/GAGAGAACCGGGGCTA-3’). Third and fourth, two phase variable O1-antigen biosynthesis genes, *manA* and *wbeL*, were locked into their ‘on’ configurations by placing silent A-to-G mutations within their poly-A tracts. This was done to reduce the frequency of appearance of phase-off mutants that no longer make O1-antigen, which is the receptor for vibriophages ICP1 and ICP3. For *manA*, which has two poly-A tracts, mutations A213G, A216G, A609G, and A612G were made. For *wbeL*, which has one poly-A tract, mutations A111G and A114G were made. Finally, the streptomycin-resistance point mutation within *rpsL* was reverted to the wild type sequence. AC6169 was subjected to whole-genome sequencing and the desired mutations confirmed.

To prepare freshly growing *V. cholerae* AC6169, the bacterial preparation starts two days in advance. From an -80°C stock vial of AC6169, we streaked a Luria-Bertani agar (LA) plate with the bacteria and incubated it overnight at 37°C. The next day, using a single colony, we inoculated a 14-ml culture tube (Falcon cat# 352059) with 5 ml of Luria-Bertani broth (LB) and incubated the tube for 14-16 hours in a shaker incubator at 37°C. From this culture we measured the OD_600_ and transferred a small volume of the overnight culture broth into a tube with 5 ml LB broth to prepare a broth with an OD_600_ of 0.05 in the new tube. We then incubated the new tube with shaking at 37°C to reach an OD_600_ of 0.2.

We prepared a 0.06% solution of resazurin (Invitrogen, cat#R12204) by dissolving it in autoclaved milli-Q water in a 50 ml tube. We vortexed the mixture until the resazurin was completely dissolved. We then filtered it through a 0.22 µm filter (Millipore, cat#SLGVR33RS). We prepared aliquots (550 µl) in 1.5 ml sterile tubes and stored them at -20°C in a foil lined box. When needed, (within 30-60 minutes), we removed an aliquot(s), covered them with foil, and thawed them at room temperature.

To prepare a standard lot of ICP1, we amplified it in the host strain AC6169. First, we streaked an LA plate with frozen AC6169 and incubated the plate overnight at 37°C. The next morning, we picked one colony from the LA plate and inoculated it into 5 ml LB in a 14 ml-culture tube and incubated the tube at 37°C with shaking until the OD_600_ reached 0.2-0.4. We then added 2.5 µl of ICP1 stock to the tube and continued to shake the tube at 37°C for 3 more hours. We then centrifuged the tube at room temperature at 3500 x *g* and filtered it through a 0.22 µm Millipore filter to eliminate the bacteria from the supernatant. We then stored the filtrate at 4°C until used. To quantify the titer of ICP1 in each lot, we used a standard soft agar overlay method. Briefly, we grew the host strain AC6169 to mid-log phase (OD_600_=0.3-0.5) and diluted it in LB to a OD_600_ of 0.025. We then serially diluted ICP1 in LB (10^−1^ to 10^−7^) and mixed 100 µl of the diluted AC6169 with diluted ICP1 in 96-well microplate wells in duplicate and incubated it at 37°C for 10 minutes to allow phage adsorption. We then added the total 200 µl mixture to a well with 3 ml of 48°C soft agar (0.35% agar in LB) in a 6-well plate. We gently swirled the plate immediately to mix and place the plate at room temperature until the agar is solidified. We then incubated the plate for 3-4 hours at 37°C until the plaques are observed and quantified. Based on the average number of plaques and the dilution factor, we calculated plaque titer.

### Proof of concept

As an initial experiment to establish the proof of concept, we serially (10-fold) diluted ICP1 in 5-ml culture tubes with 0.25 ml LB using concentrations of 4 to 400 pfu/ml. We then added 250 µl fresh LB and 35 µl of freshly growing AC6169 which had been adjusted to an OD_600_ of 0.2 and incubated the mixture with shaking at 37°C for 2, 2 ½ or 3 hours. After the incubation, we added 12.5 µl of the resazurin solution (0.06%) and continued the shake incubation for another 30 minutes. We then centrifuged the tubes (3650 x *g)* and observed the tubes visually to assess their color and photographed the tubes to record their color. To obtain an objective measure of the color change, we diluted 50 µl of the supernate in 950 µl LB broth and measured the OD_600_. The OD_600_ of each sample and the controls were recorded and the result was expressed as the ratio of the sample OD_600_ divided by the OD_600_ of the negative control sample (without phage). We expect the negative control sample to have a low OD_600_ and samples with phage should have a significantly higher OD_600_ than the negative control, with a ratio >2.

### Procedure to detect phage in test samples

To detect phage in different sample types spiked with ICP1 (bay water, stool, and bile peptone broth), we evaluated two methods. One method, we call the one-step method, we only amplify the phage once after the sample is made bacteria free. The other method we call the two-step method since we amplify the phage twice.

For the one-step method, for each sample, we centrifuged the sample for 5 minutes at 17,000 *x g* at room temperature to rid the sample of debris. We then prepared a bacteria-free sample by filtering the supernate through a 0.22 Millipore filter or by treating it with chloroform. When we used chloroform, we transferred 1 ml of the supernate to a new 1.5 ml tube and added 100 µl of fresh chloroform and vortexed it for 15 seconds. We then centrifuged it at room temperature for 5 minutes at 17,000 *x g* and then transferred 600 µl from the top of the tube to a new sterile 1.5 ml centrifuge tube. We centrifuged it again for 5 minutes at 17,000 *x g* and we transferred 250 µl from the top of the tube to a new sterile 5 ml culture tube to be used as the bacteria-free sample for further testing in the same manner as the filtered sample.

Using the bacteria-free sample we proceeded to detect the phage. Using a separate 5 ml culture tube, we mixed 250 µl of the bacteria-free sample with 250 µl LB and 35 µl of freshly growing AC6169 which had been adjusted to an OD_600_ of 0.2 and incubated the mixture with shaking at 37°C for three hours. After the 3-hour incubation, we added 12.5 µl of the resazurin solution (0.06%) and continued the shake incubation for another 30 minutes. We then centrifuged the tubes for 10 minutes at 3650 x g and observed the tubes visually to assess their color and photographed the tubes to record their color. As was done in the pilot experiment, to obtain an objective measure of the color, we diluted 50 µl of the supernate in 950 µl LB broth and measured the OD_600_. The OD_600_ of each sample and the controls were recorded and the final result was expressed as the ratio of the sample OD_600_ divided by the OD_600_ of the negative control sample (without phage). The negative control sample will have a low OD_600_ and those with phage should have a significantly higher OD_600_ than the negative control, generally the ratio will be >2.

For the two-step method we amplified the phage as the first step without resazurin. To amplify the phage, we centrifuged the sample for 5 minutes at 17,000 *x g* at room temperature to rid the sample of debris and mixed 750 µl of the supernate with 675 µl LB and 75 µl of freshly growing *V. cholerae* AC6169 (OD_600_ adjusted to 0.2) in a 5 ml culture tube. We then incubated the culture tubes while shaking for 2 hours at 37°C. We then centrifuged the tube at room temperature for 5 minutes at 17,000 x *g*. We then collected 1 ml of the supernate and rendered it bacteria-free, either by filter sterilizing through a 0.22 µm Millipore filter or by treating with chloroform as described above.

After amplifying the phage, as a second step, we proceeded to detect the phage as was done similarly in the one-step method except that the incubation was for two hours rather than three as in the one-step method before adding the resazurin. The two-step procedure can be done in one day, but it can also be spread over two days by storing the bacteria-free, phage-amplified sample overnight at 4°C. Both the initial amplification (first step) and the detection (second step) need fresh AC6169. A schematic of the procedures for the two-step method is shown in Figure 1.

**Figure 1.**
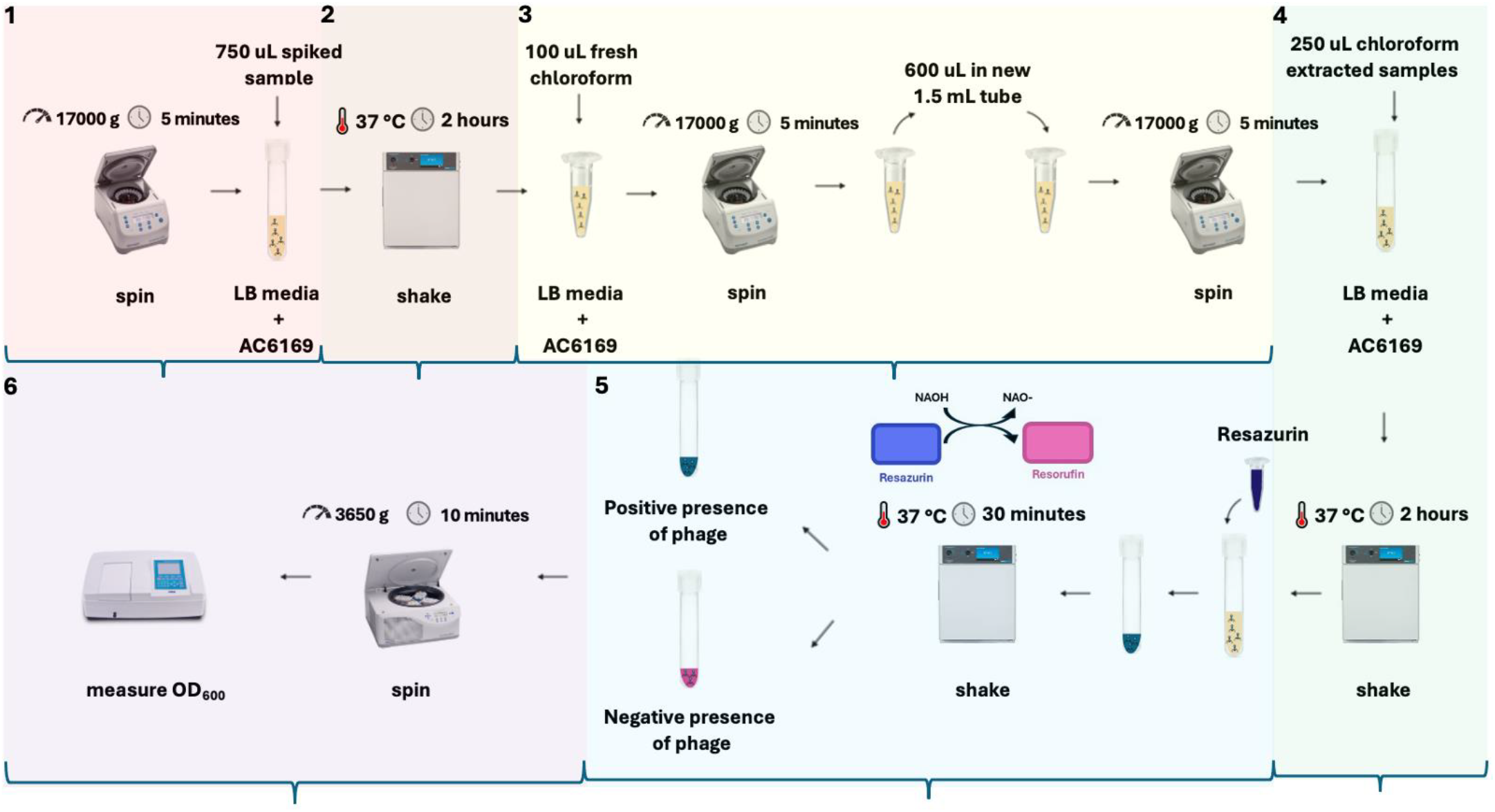
Schematic illustrating the two-step amplification procedure for detecting vibriophage. The steps are indicated by the numbers in each panel. 1) centrifugation of sample and collection of supernate, 2) shaking incubation of the supernate with LB and AC6169 for two hours, 3) centrifuge and preparation of bacteria-free supernate using chloroform (centrifuge 2X), 4) shake-incubation of the bacteria-free sample with LB and AC6169 for two hours, 5) addition of resazurin and continue shaking incubation for 30 additional minutes, 6) centrifuge culture and observe color change of supernate. Photograph and measure OD_600_.

To determine the sensitivity of the assay, we spiked the water, bile peptone or stool sample with different concentrations of ICP1 ranging from 4 to 400 plaque forming units (pfu) per ml. For bile peptone broth, after spiking the sample, we froze it at -80°C and later we thawed the sample for testing. For each assay, we included a negative control with AC6169 but without phage, and a second control with neither phage nor AC6169 (dye only control)

## Results

The results of the proof-of-concept experiment are shown in Figure 2. A clear dose response was observed when the initial incubation was limited to 2 hours, but with longer times of incubation, the bacteria continued to grow and the blue color in each of the tubes increased, obscuring the dose response.

**Figure 2.**
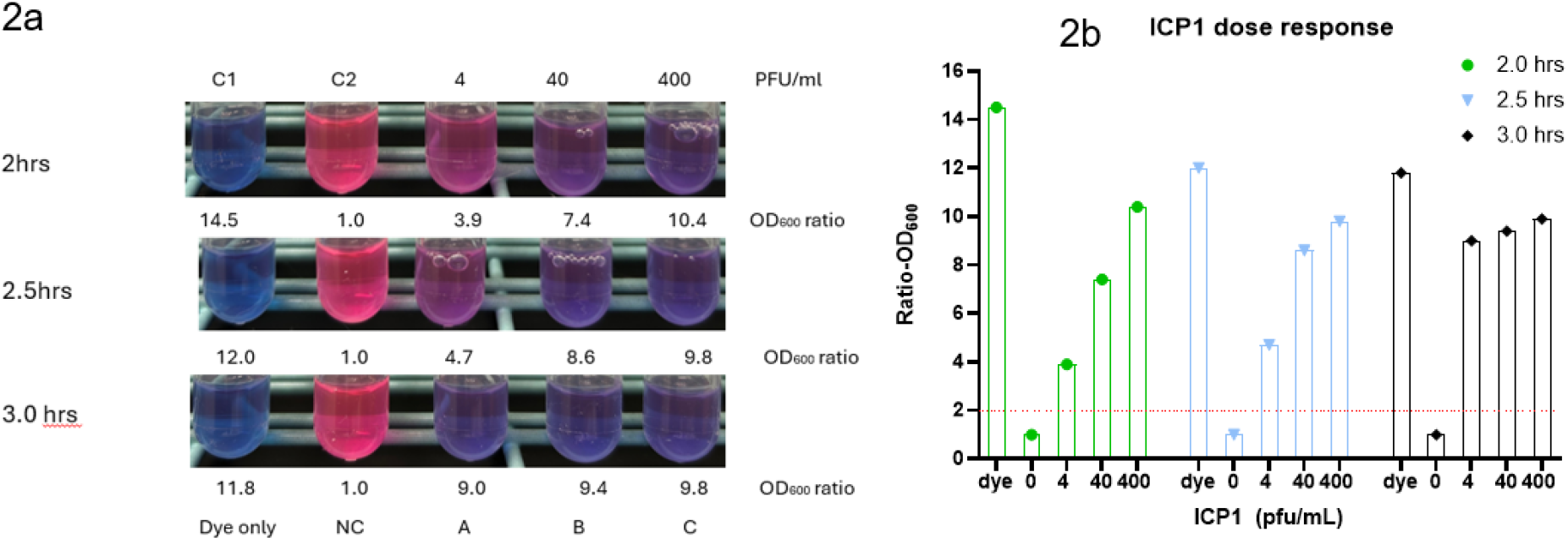
2a: Color reactions in tubes using serial dilutions of phage in LB broth with AC6169. Control sample (C1) had no phage and no bacteria. Control sample (C2) is the negative control with no phage but with AC6169 bacteria. Test samples were inoculated with serial dilutions of ICP1 with AC6169 from 4 to 400 pfu /ml (A,B,C). The numbers below each tube are the ratio of OD_6oo_ of the test sample compared to the negative control (NC). 2b: Shows these results graphically.

To determine if the assay can detect phages in different sample types using the one-step method, we spiked samples of stool, bay water, and bile peptone with serial dilutions of ICP1 and tested the filtrates from a 0.22 Millipore filter or chloroform extraction. Figure 3 shows the OD ratios of the experiments with the different sample types using either Millipore filtration (3a) or chloroform (3b). The limit of detection for these sample types was from 4 to 40 pfu/ml for the Millipore filtration but was less sensitive using the chloroform procedure (40 to 400 pfu/ml).

**Figure 3.**
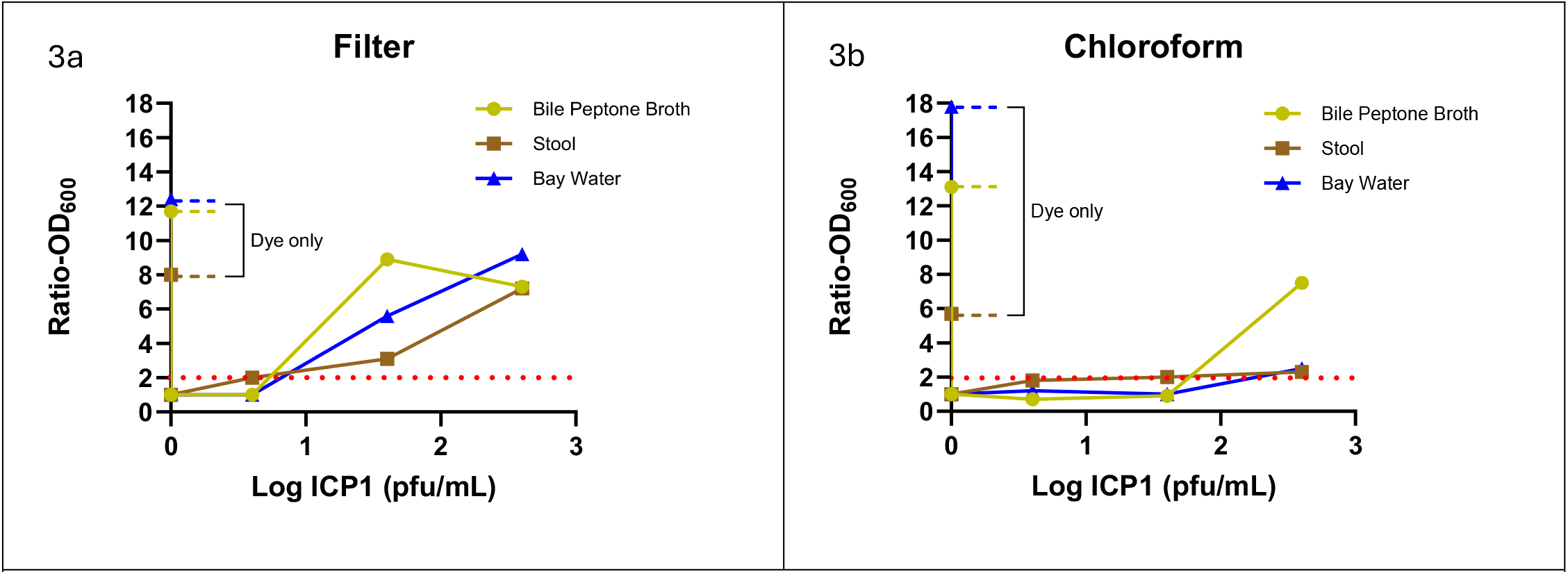
shows the OD ratios of the different sample types using the one-step method. 3a used Millipore filtration and 3b used chloroform to prepare the bacteria-free sample. The red dotted line shows a ratio of 2.0 above which the sample is considered positive.

As shown in Figure 4, we found that prior amplification of phages through a two-step procedure improved the sensitivity of the assay when using chloroform for all three sample types. The limit of detection for these samples was from 4 to 40 pfu/ml.

**Figure 4.**
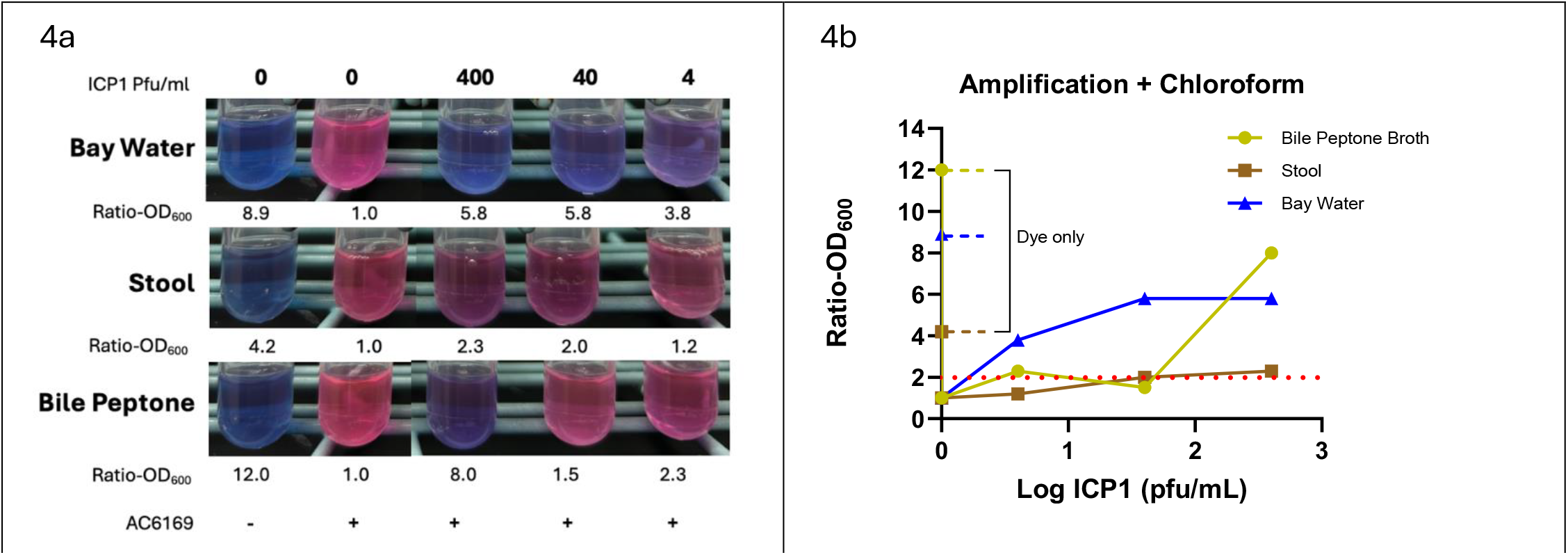
4a shows the colors of the tubes when ICP1 was serially diluted in samples of bay water, stool, and bile peptone broth and incubated with AC6169 and resazurin. The ratio of OD_600_ was the value from each tube divided by the negative control without ICP1. The numbers below each tube are the ratio of OD_600_ of the test sample compared to the negative control (NC). 4b shows the ratios graphically. The red dotted line shows a ratio of 2.0 above which the sample is considered positive.

## Discussion

The study findings show that it is possible to screen samples of environmental water, stool, and frozen bile peptone broth for virulent vibriophages (phage) using a method that leverages a dye (resazurin) that in the experimental conditions detects bacterial lysis. When the bacterium is allowed to grow, in the presence of resazurin, the color of the broth changes from blue to pink; however, if vibriophages are present, the phage lyse the bacteria, the bacteria cannot activate the dye, and the color remains blue. This assay uses *V. cholerae* AC6169 to amplify and detect the phage because this strain is receptive to phage specific to *V. cholerae* (ICP1, 2, and 3). Generally, we are able to discern positive samples by comparing the colors of the tubes using the naked eye; however, we found that some samples are not as clear and measuring the OD_600_ provides a more objective outcome.

In the assay, AC6169 is the only agent in the test condition to convert resazurin. To minimize false positives, the assay requires that the test sample must be devoid of other biologic agents (bacteria) and inorganic sources of NADH oxidase.

The bacteria in the test sample can be eliminated either passing the sample through a 0.22 µ filter or by killing the bacteria with chloroform. For each sample type, we confirmed that the filtered (or chloroform extracted) samples were free of contaminating bacteria by adding equal volume of LB broth and grow at 37°C overnight. While filtration may be gold standard, the filters are costly. The cost of a one-step procedure per sample was estimated at $4 using filtration and only $1.7 using chloroform. Compared to filtration, use of chloroform does require additional centrifugation steps to reduce the residual leftover of chloroform which adds to the workload.

Initially we determined if filtered water and fecal samples could be tested without the initial step of phage amplification. We assumed that a few phages would be able to attack the target bacterium AC6169 and would be detectable even without prior amplification. However, we found that the two-step assay was more sensitive when the sample is first mixed with AC6169 to amplify the phage before the sample is rendered bacteria-free for the detection step with AC6169 bacteria in LB broth and resazurin. Of note, for both the one and two step methods, we incubated the sample with LB for 2 or 3 hours before adding resazurin and then continued incubation for an addition 30 minutes.

We feel this assay shows promise for screening samples of environmental water for vibriophages, especially wastewater, for evidence of cholera transmission in the area. The bacterium *V. cholerae* can also occasionally be detected in contaminated water and this assay for phage does not negate the need for detection of the bacteria either by culture or PCR. It simply allows for phage detection as a proxy for *V. cholerae* when the bacteria are lower in number or degraded. Previously we described a rapid diagnostic test assay for detection of *V. cholerae* using samples of environmental water enriched in alkaline peptone water, and this method also provides results within a day(20). The circumstances for which assay for *V. cholerae* or vibriophage is most appropriate or most sensitive need to be determined.

We examined the recovery of phage from frozen bile peptone broth because one of our partners in Africa had been collecting Moore swabs from environmental samples to study another pathogen, *Salmonella* Typhi. This lab had saved frozen samples of Moore swabs that had been incubated in bile peptone broth before freezing. Thus, we wanted to determine if phage would survive and could be recovered from these frozen samples. We have not yet attempted to recover vibriophage from other types of frozen samples, but we found that viable *V. cholerae* O1 could not be recovered from spiked bile peptone broth.

In addition to environmental samples, we found that this assay can also be used to screen stool samples for evidence of cholera. Currently, confirmation of a cholera case depends on the detection of fecal *V. cholerae* by culture or PCR, but detection of the vibriophage in the patient’s stool also provides confirmatory evidence that the patient had cholera.

The strength of this method includes its sensitivity, its low cost, and the rapid turnaround. Results can be available within a few hours and this rapid turnaround time allows for additional samples to be collected up or downstream from the original sampling site in an attempt to better localize the source and spread of the cholera contamination.

The test does have some limitations. The one-step method using a two-hour initial incubation time may be considered to be semi-quantitative but the longer incubation times with the one step method or the two-step method are not quantitative. Secondly, we have tested these three sample types, but there may be other sample types that need to be evaluated. Thirdly, we found that the target bacteria, AC6169, was very useful but there may be other strains which may be more widely available. Fourth, we used the phage ICP1 to develop the assay and to use as a control reagent. As phages are identified in different labs, these labs may want to save their local phages to use as a control for their assay.

In terms of the next steps, we view this as a screening test for virulent vibriophage as a proxy for pathogen detection. However, the method requires further evaluation with clinical and environmental samples in regions with active cholera outbreaks. Positive samples will likely need subsequent confirmation using either PCR or plaque assays(8, 21) for phages with paired assays for *V cholerae*. Finally, while AC6169 is known to be susceptible to these three phages, there may be other vibriophages to which AC6169 is not susceptible; these phages may be detected using traditional plaque assays using other strains of *V. cholerae*. Finally, the epidemiology and ecology of virulent phages specific to *V. cholerae* needs to be further investigated and we feel these methods can be used to better understand the role that vibriophages play in transmission and the epidemiology of cholera.

## Authors contributions

WL Conceptualization, Data curation, Formal analysis, Investigation, Methodology, Supervision, Writing review and editing

DD Data Curation, Formal Analysis, Investigation, Methodology, Visualization, Writing – Review & Editing AC Funding Acquisition, Investigation, Resources, Writing – Review & Editing

JRA Conceptualization, Funding Acquisition, Methodology, Writing – Review & Editing KEdaS Conceptualization, Methodology Writing – Review & Editing

AKD Conceptualization, Project Administration, Writing – Review & Editing MAS Investigation, Resources

EJN Conceptualization, Investigation Resources, Supervision Writing – Review & Editing

DAS Conceptualization, Formal Analysis Funding Acquisition Investigation Methodology Project Administration Resources, Supervision Writing – Original Draft Preparation

